# An Efficiency Targeting Parameter Space for Personalized 4×1 HD-tES: Montage Description, Optimization and Application

**DOI:** 10.64898/2026.03.25.714169

**Authors:** Farui Liu, Shuhao Luo, Kexian Wang, Yuanyuan Chen, Zeqing Zheng, Haosen Cai, Tianshu Chu, Chaozhe Zhu

## Abstract

**Background:** Personalized optimization of 4×1 high-definition transcranial electrical stimulation (HD-tES) faces inherent trade-offs between montage flexibility, computational efficiency, and implementation accessibility. Conventional 10-10 electrode systems constrain placement to discrete landmark positions, while unconstrained optimization relies on stochastic algorithms that risk converging to local optima and requires neuronavigation equipment often unavailable in rehabilitation settings. Here we introduce a scalp geometry-based parameter space (SGP) that parameterizes 4×1 HD-tES montages using three intuitive scalp-defined parameters—position, radius, and orientation—and characterize parameter-performance regularities through exhaustive electric field simulations across 30 subjects and 624 cortical targets (>3.6 million configurations).

**Results:** Position primarily determined proximity to optimal performance, radius governed the intensity-focality trade-off, and orientation served as fine-tuning. Exploiting these regularities, a minimal search space (SGP-MSS) was constructed that reduced computational complexity by over 90% while guaranteeing global optima identification. Compared with standard 10-10 montages, SGP-MSS achieved up to 99% higher targeting intensity and 126% higher focality (all p < 0.0001). Compared with lead-field-free optimization, SGP-MSS achieved comparable performance with greater cross-subject stability.

**Conclusions:** The SGP framework enables efficient individualized HD-tES optimization without neuronavigation. Its scalp-based parameterization supports electrode positioning via standard cranial landmark measurements, facilitating translation to routine clinical and home-based rehabilitation settings.

## Background

Non-invasive brain stimulation has emerged as a transformative tool for neuroscience and clinical intervention [1,2]. Among these techniques, transcranial electrical stimulation (tES) has attracted particular interest due to its low cost, portability, and favorable safety profile, enabling applications ranging from laboratory research to home-based therapy [3–5]. However, stimulation effects vary considerably across individuals, and the complexity of findings across studies has made interpretation challenging [6,7]. This variability arises from multiple sources of individual differences, including neuroanatomy, functional organization, and brain network architecture [8–10]. Anatomical variations produce substantial differences in electric field distributions even under identical electrode configurations, while functional and network-level differences further influence how individuals respond to a given stimulation pattern. These observations have motivated the development of individualized optimization approaches, in which electrode placement is tailored to each participant’s specific characteristics [11–13].

High-definition transcranial electrical stimulation (HD-tES), particularly the 4×1 ring configuration, represents a significant advance toward more focal and targeted stimulation [14–16]. By positioning four return electrodes concentrically around a central electrode, 4×1 HD-tES generates a unimodal electric field with spatial focality approaching that of transcranial magnetic stimulation (TMS) [17]. This configuration exhibits a fundamental intensity-focality trade-off: larger electrode radii yield higher stimulation intensity but reduced spatial focality, whereas smaller radii enhance focality at the cost of intensity [16]. The enhanced precision of 4×1 HD-tES makes individualized optimization particularly valuable—yet also more challenging, as optimal electrode placement requires careful consideration of both the target location and the desired balance between intensity and focality.

Current approaches to 4×1 HD-tES optimization face inherent trade-offs among montage flexibility, computational efficiency, and implementation accessibility. In practice, electrodes are typically placed according to the 10-10 EEG system for operational convenience [18–20]. However, this discrete landmark system constrains electrode positions to predetermined locations, preventing precise targeting of cortical regions that fall between landmarks and limiting the flexibility to adjust electrode radius for an optimal intensity-focality balance. To overcome these constraints, recent work proposed continuous parameterization schemes using ellipsoid-based coordinate systems combined with lead-field-free optimization [21]. While this approach enables flexible electrode placement, it relies on stochastic optimization algorithms (e.g., differential evolution) that may converge to local optima rather than identifying globally optimal configurations. Furthermore, implementation requires neuronavigation systems for electrode positioning, limiting accessibility in routine clinical settings and home-based applications where such equipment is unavailable. This constraint is particularly relevant for neurorehabilitation, where multi-session protocols demand practical, repeatable electrode placement that patients or caregivers can perform independently.

A key observation motivates the present work. Because 4×1 HD-tES generates a unimodal field distribution centered beneath the central electrode [16]—unlike modalities that produce complex, multi-focal patterns (such as temporal interference stimulation)—its parameter-performance relationships may exhibit exploitable structure. If so, a constrained search space can be constructed that dramatically reduces computational complexity while guaranteeing global optima identification, a property that stochastic methods cannot provide. Furthermore, if electrode configurations are parameterized directly on the scalp surface rather than in an abstract coordinate system, the resulting montages can be implemented using anatomical landmarks and simple geodesic measurements without neuronavigation, extending individualized optimization to resource-limited clinical and home-based rehabilitation settings.

Here, we introduce a Scalp Geometry-based Parameter Space for 4×1 HD-tES (SGP-4×1 HD-tES) that addresses these challenges. The framework parameterizes electrode configurations using three intuitive scalp-based parameters—position, radius, and orientation—enabling direct specification without geometric transformation. Through comprehensive simulations across 30 subjects (over 3.6 million configurations), we reveal structured regularities in the parameter-performance landscape: position primarily determines proximity to optimal performance, radius governs the intensity-focality trade-off, and orientation serves as a fine-tuning parameter. Exploiting these regularities, we construct a minimal search space (SGP-MSS) that reduces computational complexity by over 90% while mathematically guaranteeing global optima identification. Validation against conventional 10-10 placement and state-of-the-art lead-field-free optimization demonstrates substantial improvements in targeting performance with greater stability across subjects. This strategy of characterizing parameter-performance structures to guide optimization may generalize beyond 4×1 HD-tES to other neuromodulation techniques constrained by physical principles, and offers a practical pathway toward implementing individualized brain stimulation in rehabilitation and home-based settings.

## Method

### • The SGP-4×1 HD-tES Parameter Space

The SGP-4×1 HD-tES parameter space represents each electrode configuration using three intuitive, scalp-based parameters: **position (s)** of the central electrode, **radius (r)** between the central and four return electrodes, and **orientation (**φ) of the return-electrode array (Figure 1a).

**Figure 1.**
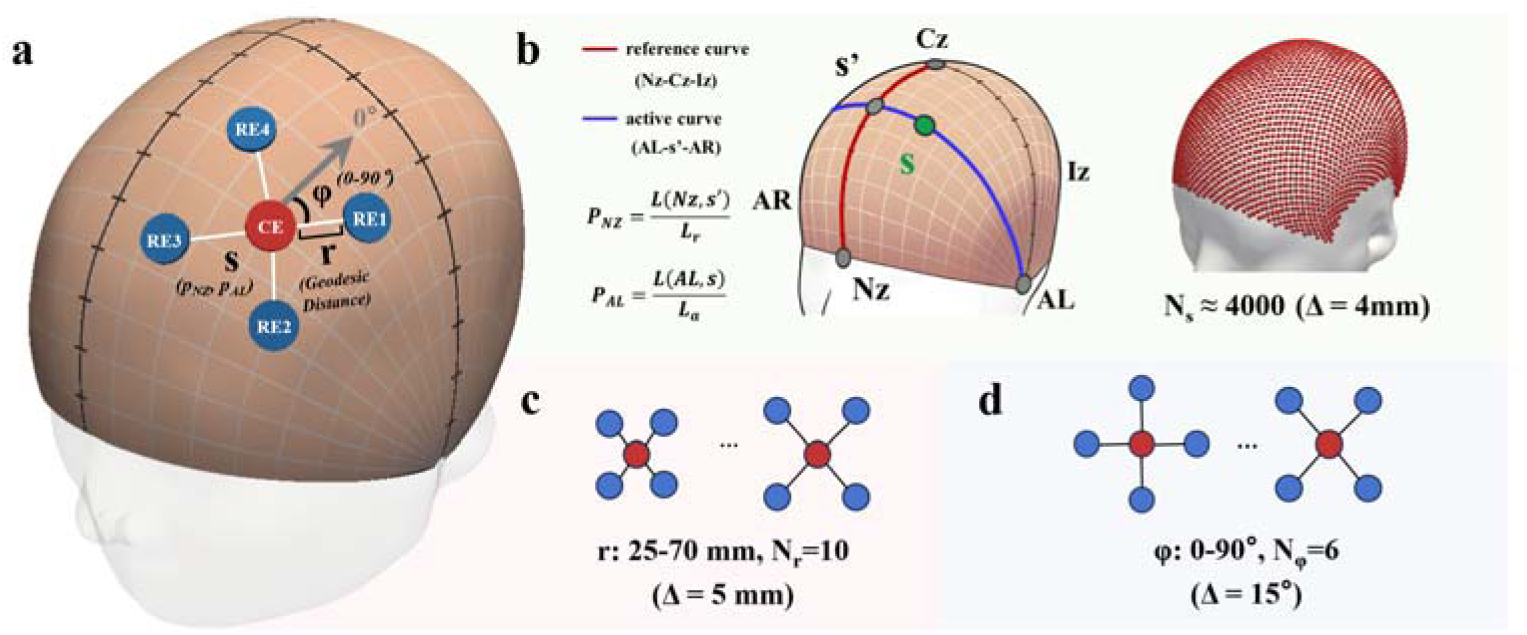
The SGP-4×1 HD-tES parameter space. **(a)** Parameterization scheme for the 4×1 HD-tES montage. The central electrode (CE, red) and four return electrodes (RE1–RE4, blue) are specified by three scalp-based parameters: position (s), radius (r), and orientation (φ). **(b)** Continuous Proportional Coordinate (CPC) system used to represent the CE position. Left: CPC is defined by five cranial landmarks (Nz, Iz, Cz, AL, AR). For any scalp site s, its coordinates (p_NZ_, p_AL_) are derived from geodesic distances along the Nz–Cz–Iz reference curve and the AL–s–AR active curve, where s′ denotes their intersection point. Right: Sampled scalp grid showing ∼4,000 valid positions (after removing periorbital and periauricular regions), with an effective spacing of ∼4 mm. **(c)** Discretization of the radius parameter (r), with ten values from 25 to 70 mm in 5-mm increments. **(d)** Discretization of the orientation parameter (φ), with six angles from 0° to 75° in 15° increments; the 0°–90° range is sufficient due to the four-fold rotational symmetry of the array.

#### Parameter Definition

**Position (s)**:The central electrode position is specified using the Continuous Proportional Coordinate (CPC) system [22–25], a standardized scalp-coordinate system defined by five cranial landmarks: nasion (Nz), inion (Iz), vertex (Cz), and left and right preauricular points (AL, AR). For any scalp location, its anterior-posterior coordinate (p_NZ_) is computed as the normalized geodesic distance along the Nz-Cz-Iz reference curve, while its left-right coordinate (p_AL_) is computed along the coronal curve passing through that location (Figure 1b, left). Both coordinates range from 0 to 1, enabling continuous parameterization with cross-subject anatomical correspondence.

**Radius (r):** The radius is defined as the scalp geodesic distance between the central electrode and each of the four return electrodes (Figure 1c).

**Orientation (**φ): Orientation is defined as the clockwise angle from the posterior direction to the first return electrode, ranging from 0° to 90° due to four-fold rotational symmetry (Figure 1d).

#### Parameter Space Discretization and Feasibility Constraints

Candidate positions were sampled uniformly across the scalp with approximately 4 mm spacing (see Supplementary Material for details). Regions unsuitable for electrode placement, including periorbital and periauricular areas, were excluded, yielding approximately 4,000 valid central electrode positions (Figure 1b, right). Radius was discretized into 10 values (25–70 mm, 5 mm increments), covering the commonly used range for 4×1 HD-tES. Orientation was discretized into 6 values (0°–75°, 15° increments), which is adequate given the array’s four-fold rotational symmetry. For each valid position, only radius-orientation pairs that keep all four return electrodes within suitable scalp regions were retained. This procedure yielded approximately 120,000 feasible parameter combinations per subject (see Supplementary Material for detailed coverage analysis).

### • Targeting Performance Space

Each parameter combination (s, r, φ) was evaluated using two complementary metrics within a Pareto optimization framework [26,27].

**Targeting intensity (*I*)** is defined as the volume-weighted average electric field magnitude within the region of interest (ROI):

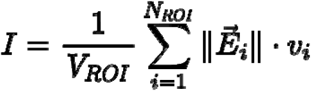

Where 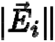 is the electric field magnitude in the ***i***-th mesh element within the ROI,***v_i_*** is the volume of that element,***V_ROI_*** is the total volume of the target region, and ***N_ROI_*** is the total number of mesh elements within the ROI. This metric reflects the strength of stimulation delivered to the target.

**Targeting focality (*F*)** is defined as the ratio of the target-region average field magnitude to the gray-matter average:

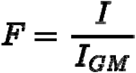

where ***I_GM_*** is the volume-weighted average electric field across cortical gray matter. Higher values indicate greater spatial specificity of stimulation.

All feasible parameter combinations (∼120,000 per subject) were exhaustively evaluated. A configuration was considered Pareto-optimal if no other configuration achieved both higher intensity and higher focality simultaneously. The resulting Pareto front represents the set of optimal trade-offs between these competing objectives.

### • Parameter-Performance Relationship in SGP-4×1 HD-tES

To investigate how the three SGP parameters shape stimulation performance, comprehensive electric field simulations were conducted across the full parameter space for each individual head model.

For position, the nearest scalp point ***s_0_*** was first identified as the scalp location with minimal perpendicular distance to the ROI center. The geodesic distance ***d(s, s_0_)*** between each candidate position s and ***s_0_*** was then computed, and configurations were grouped by distance intervals to evaluate how proximity to the target influences performance. For radius, configurations were grouped by their radius values (25–70 mm) to examine how electrode spacing affects the intensity-focality trade-off. For orientation, configurations sharing the same position and radius were grouped, and an ellipse was fitted spanning the range of intensity and focality across the six orientation values. The ellipse area served as a summary measure of orientation-related variability.

### • Participants and MRI Data

Thirty healthy participants (15 females, 15 males; mean age = 20.3 ± 1.31 years) from the Southwest University Longitudinal Imaging Multimodal (SLIM) dataset were included [28]. Data collection was approved by the Research Ethics Committee of the Brain Imaging Center at Southwest University (approval number: spy-2012–012; June 3, 2012). High-resolution T1-weighted structural MRI scans (1 × 1 × 1 mm³) were acquired using a magnetization-prepared rapid gradient-echo (MPRAGE) sequence (see Supplementary Material for detailed acquisition parameters).

### • Volume Conductor Model Construction

Individual volume conductor models were constructed using the *headreco* pipeline in SimNIBS 3.2.6 [29], which performs automated tissue segmentation and generates tetrahedral meshes from T1-weighted MRI data. All segmentation outputs were visually inspected to ensure anatomical fidelity. The resulting models were converted to SimNIBS 4.5 format using *convert_3_to_4*. Standard tissue conductivity values were assigned (see Supplementary Material for details).

### • Electric Field Simulation

Electric field simulations were performed using the lead-field-free method in SimNIBS 4.5 [21], which substantially reduces computational load compared to conventional lead-field-based approaches. Circular electrodes with 5 mm radius were modeled, with the central electrode delivering 2 mA and each return electrode delivering 0.5 mA. For each subject, approximately 120,000 configurations were simulated, requiring roughly 16 hours of computation time.

### • Target Region Definition

Four cortical regions commonly targeted in cognitive and clinical studies were selected: left primary motor cortex (L_M1, MNI: [–34, –14, 67]) [30], left dorsolateral prefrontal cortex (L_DLPFC, MNI: [–42, 44, 40]) [31], left inferior parietal lobule (L_IPL, MNI: [–47, –68, 36]) [32], and right temporoparietal junction (R_TPJ, MNI: [54, –44, 18]) [33]. MNI coordinates were transformed into each subject’s native space using nonlinear deformation fields from the *headreco* pipeline. Each ROI was defined as a 10 mm radius sphere restricted to cortical gray matter. Sensitivity analyses using 5 mm and 15 mm spheres are provided in the Supplementary Material.

### • Minimal Search Space Construction and Validation

#### • Construction Strategy

Prior work has shown that 4×1 HD-tES produces a unimodal electric field centered beneath the central electrode [14–16], suggesting that Pareto-optimal positions are likely confined to a local neighborhood around the nearest scalp point ***s_0_***. In contrast, radius and orientation may require full-range preservation as they jointly determine the focality-intensity trade-off and may vary with local cortical geometry.

To evaluate these assumptions, a whole-brain set of 624 uniformly distributed cortical ROIs was defined in MNI space, sampled at 1 cm intervals within 2.5 cm of the cortical surface following [25]. Each ROI was modeled as a 10 mm radius sphere and transformed into native space. For each subject-ROI combination (30 subjects × 624 ROIs = 18,720 combinations), exhaustive evaluation was performed to identify the Pareto front.

#### • Pareto Front and SGP Parameter Analysis

For each Pareto front, the maximum geodesic distance ***d_max_***between any Pareto-optimal position and ***s_0_*** was computed to determine the spatial extent required to capture all optimal solutions. Distributions of ***d_max_***across all subject-ROI combinations informed the neighborhood radius threshold ***d_th_*** for the SGP_MSS_. Coverage of radius and orientation values across Pareto-optimal solutions was also examined to determine whether these parameters require full-range preservation.

#### • Minimal Search Space Definition

Based on the analyses above, the SGP_MSS_ was defined as: (1) ***d(s, s_0_)*** ≤ ***d_th_***, restricting position to a local neighborhood; (2) ***r*** ∈ {25, 30, …, 70} mm, preserving the full radius range; and (3) φ ∈ {0°, 15°, …, 75°}, maintaining all orientations.

#### • Validation

The SGP_MSS_ was validated through two complementary approaches. First, under the Pareto framework, SGP_MSS_-optimized configurations were compared with conventional 10-10 montages for the four representative targets (see Supplementary Material for electrode configurations). Targeting intensity and focality were compared using paired t-tests. Second, under the ROC-based framework, SGP_MSS_ optimization was compared with both 10-10 placement and lead-field-free optimization (LFOF) implemented in SimNIBS [21]. The ROC-based metric quantifies performance as the Euclidean distance to the ideal point (0, 1) in ROC space, with smaller values indicating better performance.

#### • Statistical Analysis

Paired t-tests were used to compare targeting performance between methods. All tests were two-tailed with significance level α = 0.05. Effect sizes and exact p-values are reported throughout.

## Results

### • Targeting Performance Distribution across the SGP Parameter Space

For each subject, approximately 120,000 feasible configurations were evaluated for four representative cortical targets (L_M1, L_DLPFC, L_IPL, and R_TPJ). Figure 2 shows the distribution of targeting intensity and targeting focality for two example subjects.

**Figure 2.**
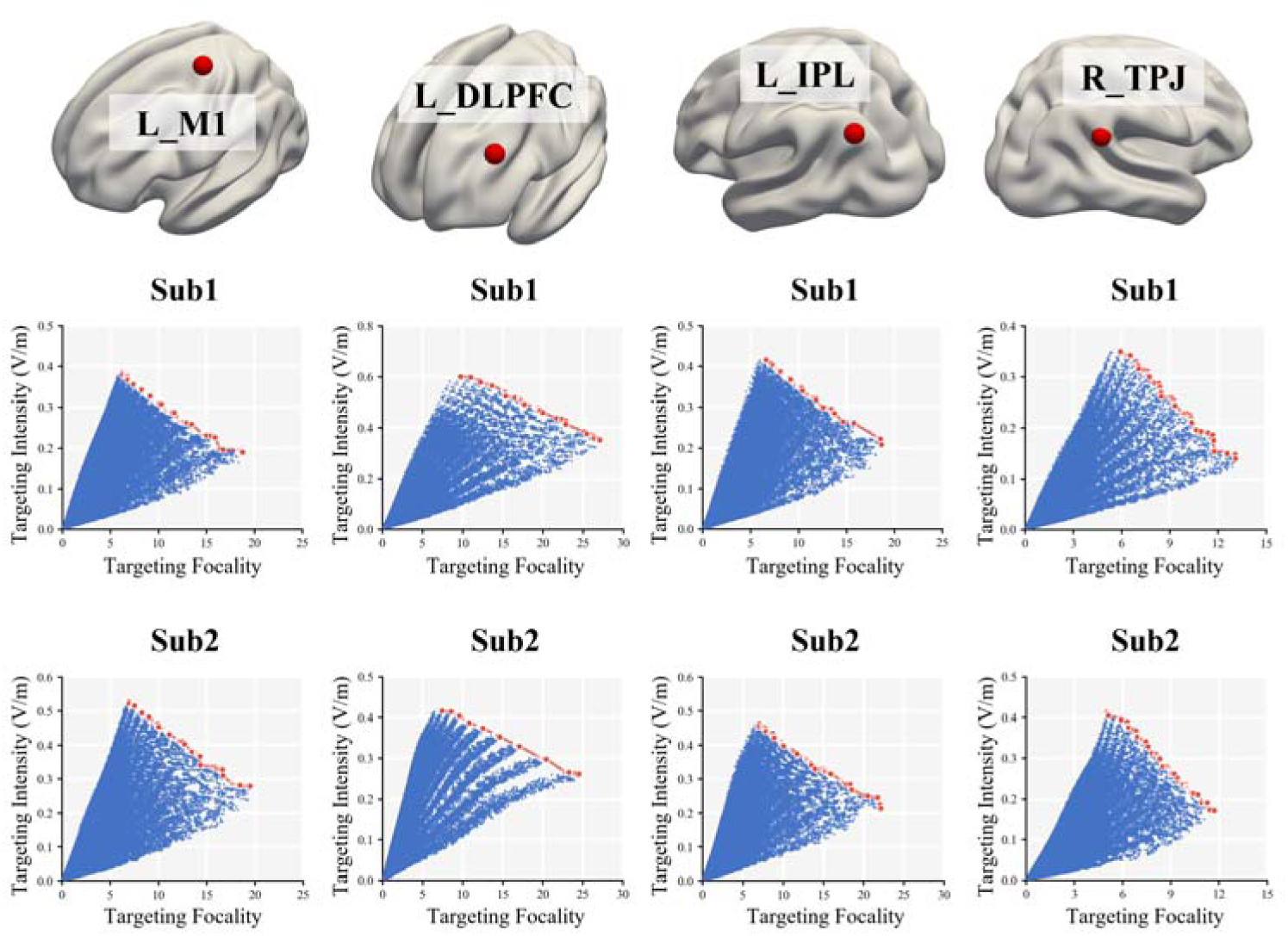
Targeting performance distribution and Pareto fronts across the SGP parameter space. Scatter plots show targeting intensity versus targeting focality for all feasible montage configurations in four representative ROIs (columns: L_M1, L_DLPFC, L_IPL, R_TPJ) from two example subjects (rows: Sub1, Sub2). Each blue point represents a single parameter combination, while red points denote Pareto-optimal solutions forming the intensity–focality trade-off boundary.

Across all targets and individuals, montage configurations exhibited a consistent performance envelope, with the Pareto front emerging as its upper-right boundary representing the best achievable trade-offs between intensity and focality. Despite substantial anatomical variability, the overall structure of this performance space remained highly stable across subjects and ROI sizes (Supplementary Material), indicating that the intensity-focality trade-off in 4×1 HD-tES follows robust modality-specific regularities.

### • Parameter-Performance Relationship in the SGP-4×1 HD-tES

Figure 3 illustrates how the three SGP parameters systematically influence targeting performance. Results are shown for the L_M1 target of a representative subject; corresponding analyses for all subjects and ROIs are provided in the Supplementary Material.

#### Effect of position (s)

Configurations were grouped by geodesic distance ***d(s, s_0_)*** between the central electrode and the nearest scalp point above the target. As shown in Figure 3a, both targeting intensity and focality decreased systematically with increasing ***d(s, s_0_)***. Pareto-optimal configurations consistently clustered within smaller ***d(s, s_0_)*** regions, whereas configurations with larger distances exhibited uniformly lower performance. Electric field visualizations (Figure 3b) confirm this distance-dependent pattern: as the central electrode moves away from ***s_0_***, target engagement becomes attenuated while off-target stimulation increases.

**Figure 3.**
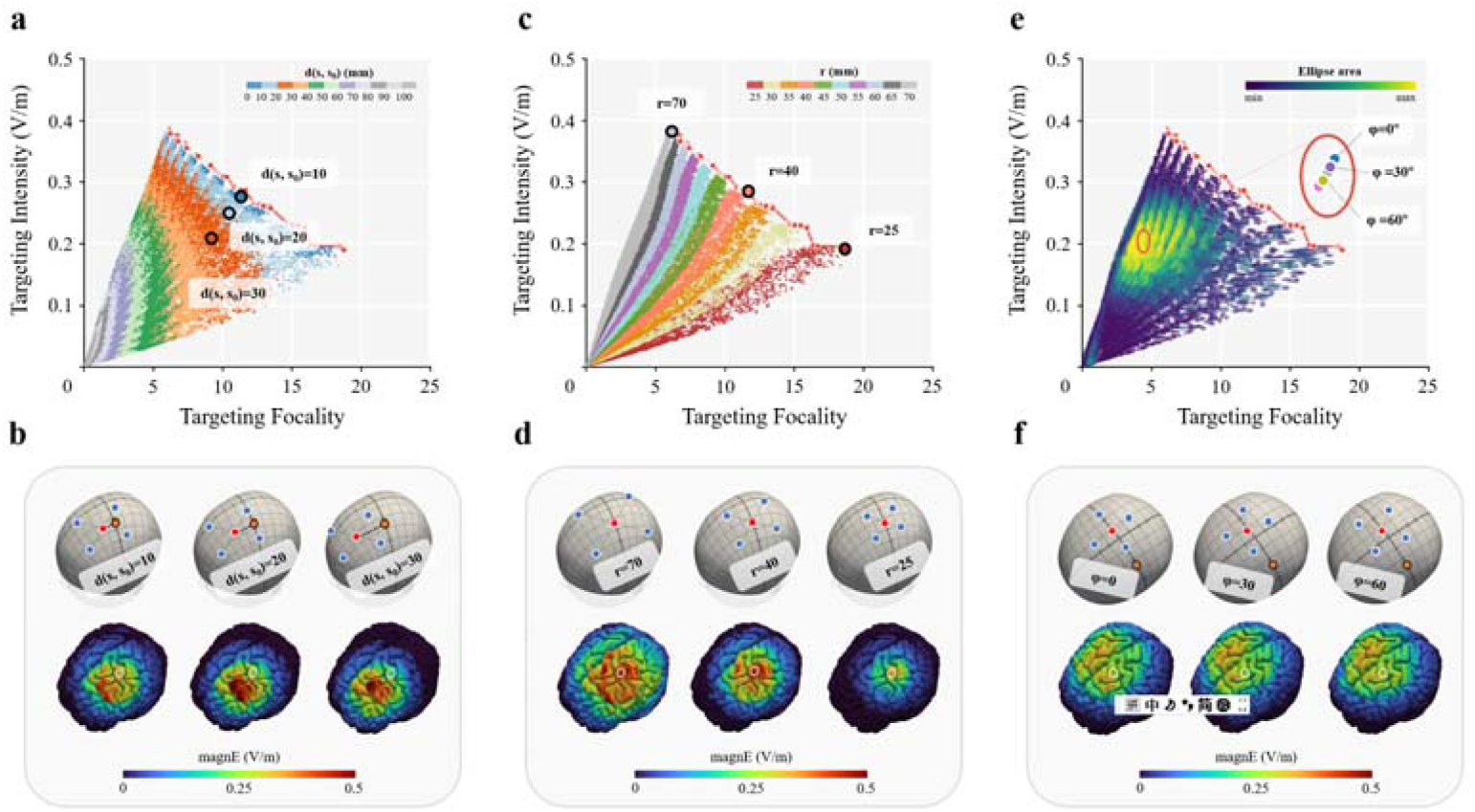
Parameter-performance relationships in SGP-4×1 HD-tES. Results are shown for the L_M1 target of a representative subject. **(a)** Distribution of targeting intensity and focality as a function of geodesic distance ***d(s, s_0_)***. Different colors represent distance intervals; Pareto-optimal configurations are highlighted. **(b)** Electric field distributions for representative configurations at varying ***d(s, s_0_)*** values (10, 20, 30 mm), demonstrating the attenuation of target engagement with increasing distance. **(c)** Distribution of targeting intensity and focality as a function of radius. Different colors represent radius values from 25 to 70 mm. **(d)** Electric field distributions for representative configurations at varying radii (r = 25, 40, 70 mm), illustrating the intensity–focality trade-off. **(e)** Distribution of targeting intensity and focality as a function of orientation. Each ellipse spans the range of intensity and focality across orientations for a fixed position–radius combination; color indicates ellipse area. **(f)** Electric field distributions for representative configurations at varying orientations (φ = 0°, 30°, 60°); configurations were selected from position–radius combinations with the largest orientation-related variability.

#### Effect of radius (r)

Varying the radius produced systematic shifts along the intensity-focality trade-off (Figure 3c). Larger radii yielded higher targeting intensity but reduced focality, while smaller radii produced more focal but weaker stimulation. Electric field visualizations (Figure 3d) illustrate this trade-off: increasing the radius expands suprathreshold stimulation extent, whereas decreasing the radius confines modulation to the target region.

#### Effect of orientation (φ)

Changes in orientation produced relatively modest effects across the parameter space (Figure 3e). For most position-radius combinations, varying φ resulted in only small shifts in performance, as reflected by predominantly small ellipse areas. Larger ellipses occurred primarily with larger radii and larger ***d(s, s_0_)***, but even in these cases variability remained limited. Electric field visualizations (Figure 3f) confirm that orientation differences in field distribution are subtle even for configurations with the largest orientation-related variability.

### • Minimal Search Space Identification

To identify the minimal parameter domain required to retain all optimal solutions, the relationship between ***d(s, s_0_)*** and performance was further examined. Figure 4a shows this analysis for the same subject and L_M1 target displayed in Figure 3. Although performance generally degrades with increasing ***d(s, s_0_)***, Pareto-optimal configurations span a broader distance range than suggested by the grouped visualization.

**Figure 4.**
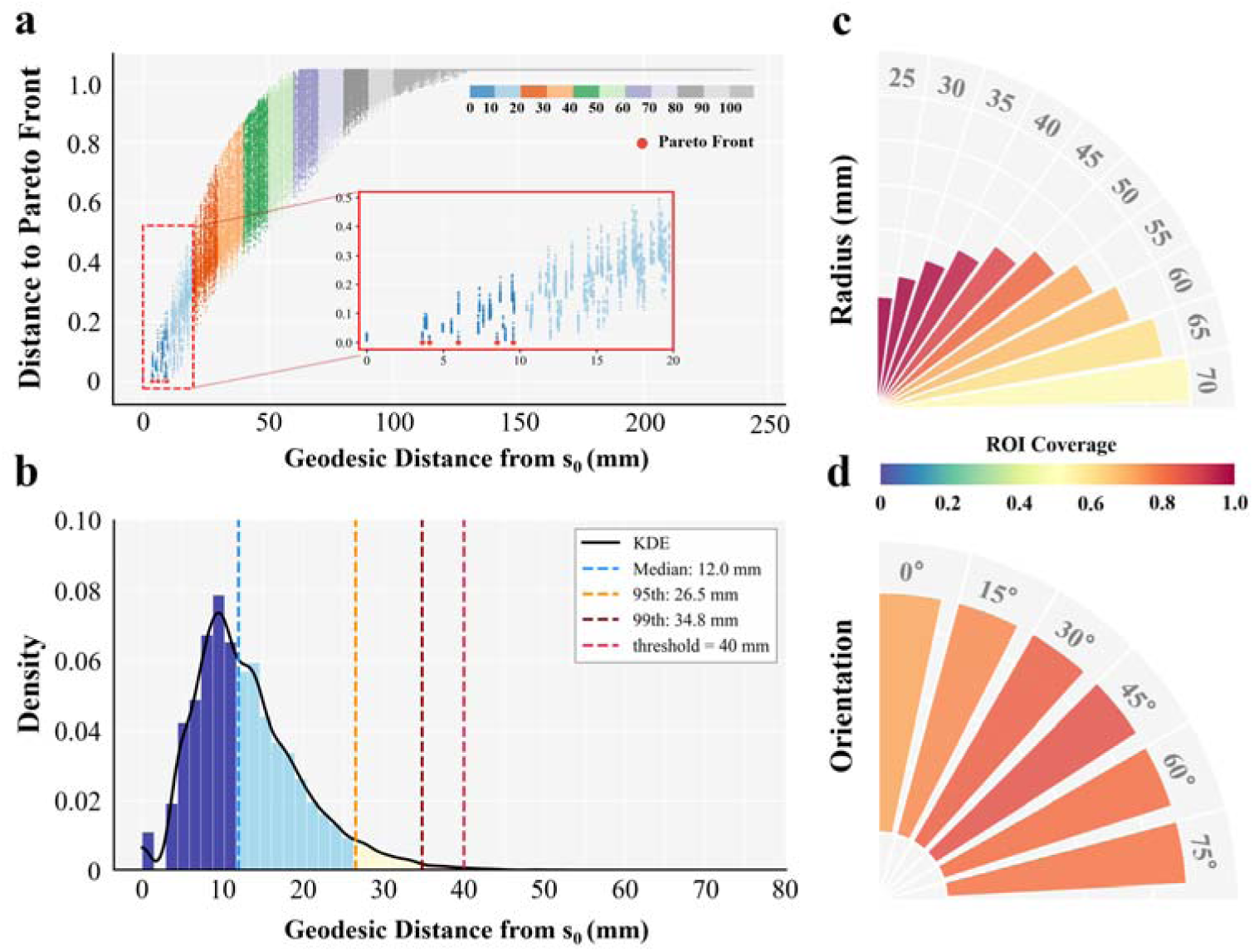
Identification of the Minimal Search Space for SGP-4×1 HD-tES. **(a)** Relationship between geodesic distance ***d(s, s_0_)*** and performance (expressed as distance to the Pareto front) for the same representative subject and L_M1 target shown in Figure 3. Although performance degrades with increasing ***d(s, s_0_)***, the inset shows that Pareto-optimal configurations span a broader distance range than suggested by the grouped visualization. Different colors represent ***d(s, s_0_)*** intervals. **(b)** Distribution of ***d_max_***—the maximum ***d(s, s_0_)*** among Pareto-optimal configurations —across all 30 subjects and 624 ROIs. Vertical lines indicate the median (12.0 mm), 95th percentile (26.5 mm), 99th percentile (34.8 mm), and the adopted threshold (***d_th_*** = 40 mm). **(c)** ROI coverage for each radius value across all Pareto-optimal configurations, showing broad coverage across all discretized values with gradual decrease at larger radii. **(d)** ROI coverage for each orientation across all Pareto-optimal configurations, with all six orientations contributing across subjects and ROIs.

To characterize this variability at the cohort level, the maximum ***d(s, s_0_)*** among Pareto-optimal configurations (***d_max_***) was computed for each subject-ROI combination. The distribution across all 30 subjects and 624 ROIs (Figure 4b) showed a median of 12.0 mm, 95th percentile of 26.5 mm, and 99th percentile of 34.8 mm. A threshold of ***d_th_*** = 40 mm was adopted to ensure complete capture of all subject- and ROI-specific variations.

The parameter ranges for radius and orientation within this neighborhood were then evaluated (Figure 4c-d). All radii contributed to Pareto-optimal sets, with coverage gradually decreasing from 99.4% at 25 mm to 51.3% at 70 mm. All six orientations were likewise represented, with no systematic exclusion observed. These findings indicate that neither radius nor orientation exhibits a restricted optimal domain, requiring full-range preservation.

Accordingly, the SGP_MSS_ was defined by d(s, s0) ≤ 40 mm while retaining all radii (25–70 mm) and orientations (0°–75°). This represents an approximately 8- to 17-fold reduction in search space, reducing computation time from approximately 16 hours to 1–2.5 hours per subject (see Supplementary Material for target-specific details), while maintaining complete coverage of Pareto-optimal configurations.

### • Validation of SGP_MSS_ Optimization

#### Validation 1: Pareto-Based Evaluation

SGP_MSS_-optimized configurations were compared with standard 10-10 montages across the four representative targets (Figure 5). Two comparison scenarios were examined: matching the focality of 10-10 placement while comparing intensity, and matching intensity while comparing focality.

**Figure 5.**
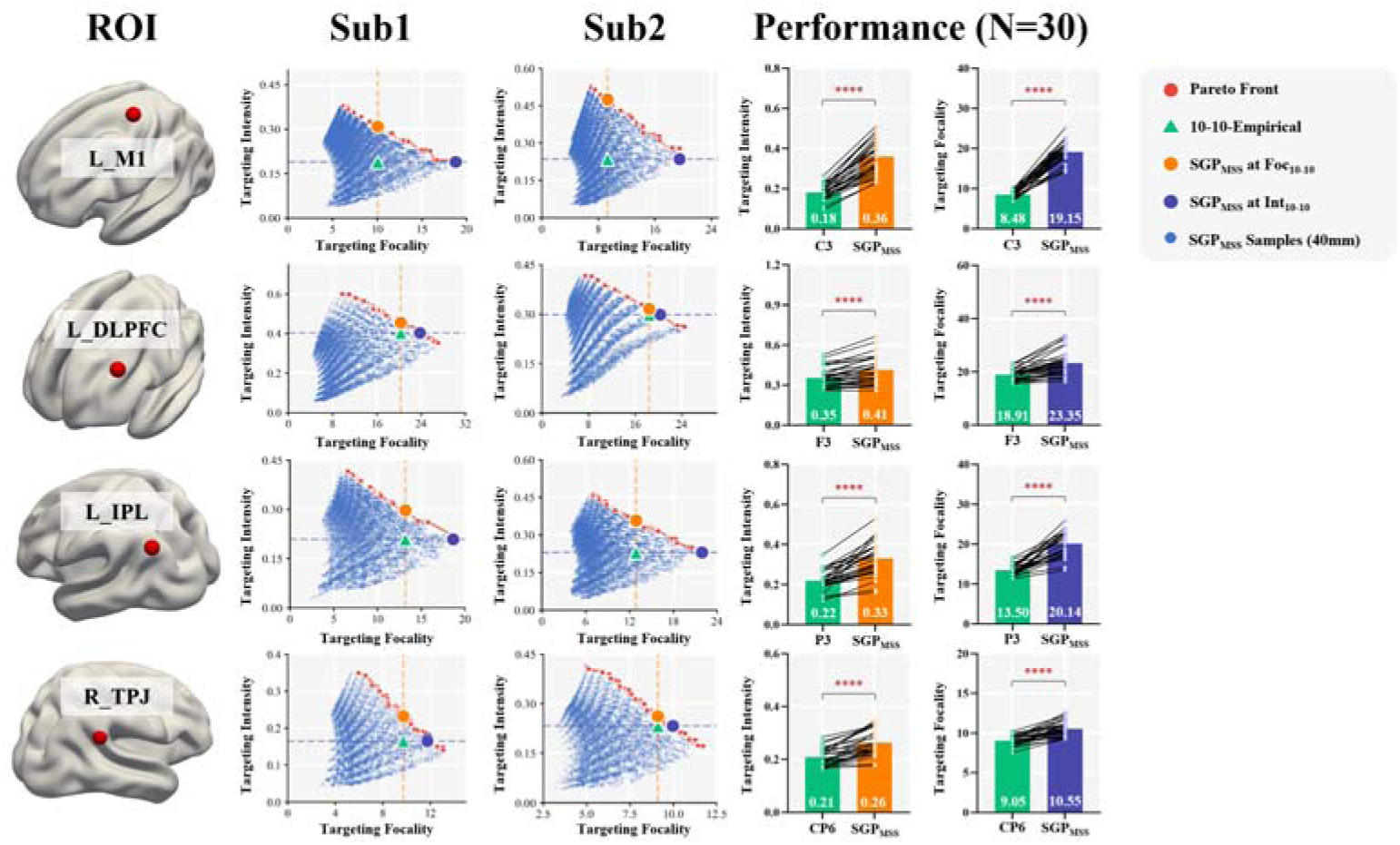
Validation of SGP_MSS_ optimization against conventional 10–10 placement under the Pareto framework. Left column: ROI locations on the cortical surface. Second and third columns: Performance distributions for two representative subjects, showing all feasible configurations (blue points), Pareto front (red points), 10–10 placement (green triangle), and SGP_MSS_ optimal configurations matched for focality (orange circle) or intensity (purple circle) of the 10–10 placement. Dashed lines indicate the focality (orange) and intensity (purple) values of the 10–10 placement. Right columns: Group-level comparison (N = 30) of targeting intensity (left) and targeting focality (right) between 10–10 placement and SGP_MSS_ optimization. Black lines connect individual subjects. Bar heights represent group means; error bars indicate standard deviation. **** p < 0.0001.

In both scenarios, SGP_MSS_ optimization substantially outperformed conventional montages. For L_M1, SGP_MSS_ achieved 99% higher targeting intensity (0.36 ± 0.08 vs 0.18 ± 0.04 V/m, p < 0.0001) when matched for focality, and 126% higher focality (19.15 ± 2.67 vs 8.48 ± 0.92, p < 0.0001) when matched for intensity. Similar improvements were observed across all other targets: intensity improvements ranged from 17% to 50%, and focality improvements from 17% to 49% (all p < 0.0001; see Supplementary Material for detailed statistics).

#### Validation 2: ROC-based evaluation

Under the ROC-based framework (Figure 6), performance was quantified as Euclidean distance to the ideal point (0, 1), with smaller values indicating better performance.

**Figure 6.**
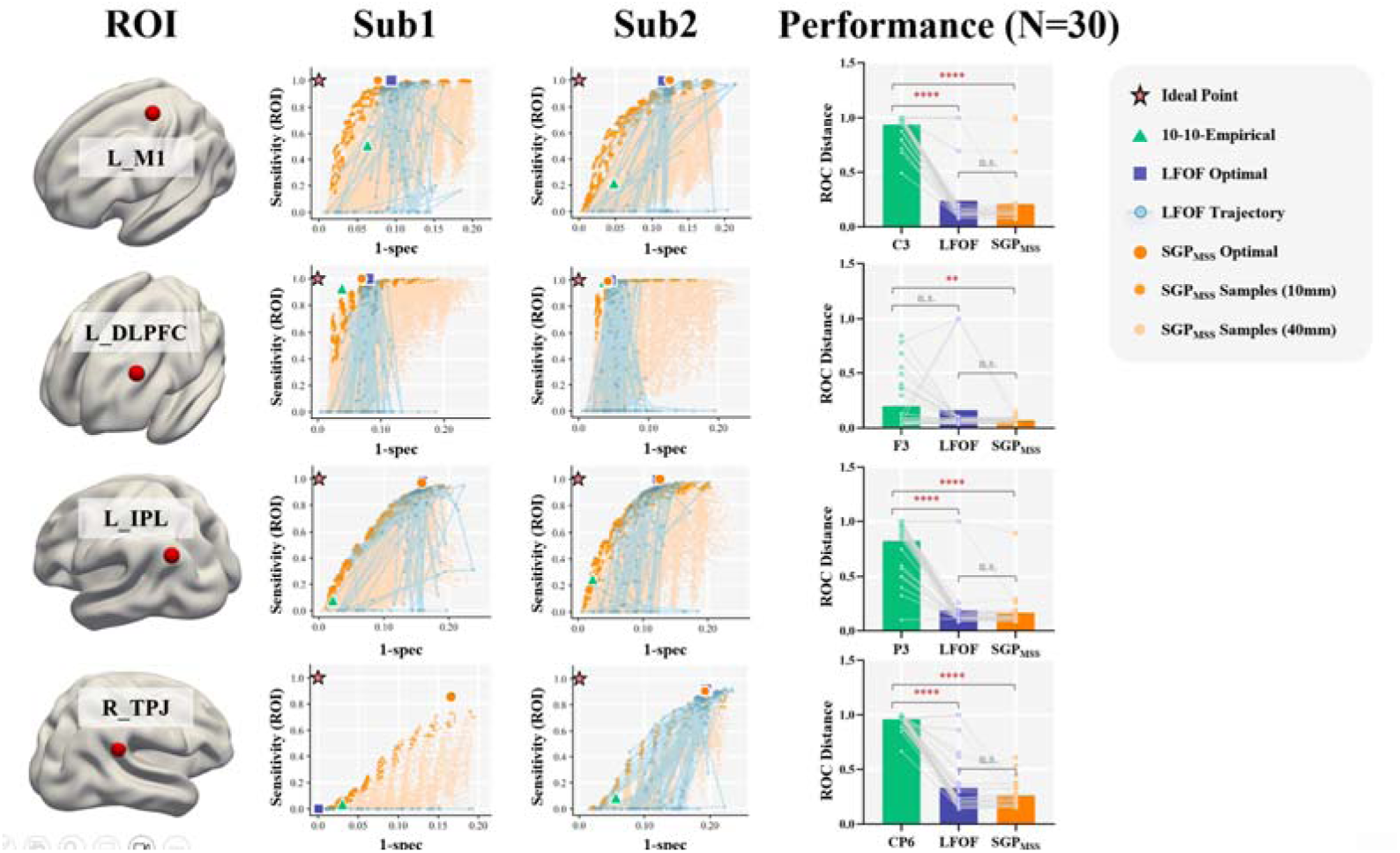
Validation of SGP_MSS_ optimization under the ROC-based evaluation framework. Left column: ROI locations on the cortical surface. Second and third columns: ROC space distributions for two representative subjects, showing SGP_MSS_ configurations (orange points, with lighter points indicating larger ***d(s, s_0_)***), LFOF optimization trajectory (blue points and lines), LFOF optimal configuration (purple square), SGP_MSS_optimal configuration (orange circle), 10–10 placement (green triangle), and ideal point (star). Right column: Group-level comparison (N = 30) of ROC distance among 10–10 placement, LFOF optimization, and SGP_MSS_ optimization. Gray lines connect individual subjects. Bar heights represent group means; error bars indicate standard deviation. **** p < 0.0001, ** p < 0.01, n.s. not significant (p > 0.05).

SGP_MSS_ optimization significantly outperformed 10-10 placement across all four targets (all p < 0.01). LFOF optimization also outperformed 10-10 placement for three targets (L_M1, L_IPL, R_TPJ; all p < 0.0001), but not for L_DLPFC (p = 0.55).

Critically, SGP_MSS_ achieved comparable performance to LFOF, with no significant differences for any target (all p > 0.05). However, LFOF converged to suboptimal plateaus in a subset of subjects (2–3 per target), resulting in substantially worse performance than SGP_MSS_ for these individuals (see Supplementary Material). Furthermore, under this single-objective framework, optimal configurations were predominantly located within ***d(s, s_0_)*** < 10 mm, suggesting that an even more constrained search space may suffice when a composite metric is employed.

## Discussion

This study introduces SGP-4×1 HD-tES, a framework that addresses the inherent trade-offs between montage flexibility, computational efficiency, and implementation accessibility in individualized brain stimulation. By characterizing parameter-performance regularities, we constructed a minimal search space that reduces computational complexity by over 90% while guaranteeing global optima identification. The framework substantially outperformed conventional 10-10 placement and achieved comparable performance to lead-field-free optimization with greater stability. When computational resources are constrained, orientation can be omitted, reducing optimization time to 10-30 minutes; for single-objective optimization, computation can be further reduced to 10-15 minutes—comparable to current SimNIBS pipelines [21].

A key methodological contribution is the systematic characterization of how montage parameters govern targeting performance, and the exploitation of these regularities to guide optimization. Conventional approaches typically treat the parameter space as an undifferentiated search domain, relying on stochastic algorithms without leveraging intrinsic parameter-performance structures [26,27]. We demonstrate that this landscape exhibits structured regularities that can be exploited to construct a minimal search space, substantially reducing computational complexity while ensuring global optima identification. This strategy may extend to other neuromodulation modalities. Although temporal interference stimulation involves more complex field patterns, systematic characterization may reveal exploitable structures. More broadly, transcranial magnetic stimulation, ultrasound stimulation, and photobiomodulation all involve parameters constrained by physical principles, suggesting similar structure-guided optimization approaches may be applicable.

Compared with existing approaches, the SGP framework offers advantages across all three stages of individualized stimulation: montage description, optimization, and implementation. The 10-10 EEG system provides straightforward electrode specification and convenient implementation with standard EEG caps, but discrete landmark positions constrain montage flexibility and limit optimization potential. The ellipsoid-based parameterization in LFOF enables continuous description, but requires geometric transformation for electrode localization, relies on stochastic algorithms that may converge to local optima, and requires neuronavigation for implementation. In contrast, SGP provides continuous parameterization with intuitive parameters defined directly on the scalp surface. By exploiting structured parameter-performance regularities, the search space permits exhaustive evaluation, guaranteeing global optima identification. Critically, scalp-based parameters enable manual electrode placement without neuronavigation, facilitating implementation in resource-limited clinical settings and home-based therapy applications.

The Pareto front identified through SGP_MSS_ optimization provides clinicians and researchers with a spectrum of optimal solutions rather than a single configuration. When maximizing stimulation intensity is the priority, configurations at the high-intensity end can be selected; when avoiding off-target stimulation is critical, as in cognitive neuroscience studies requiring precise functional attribution, high-focality configurations are preferable [34]. The Pareto framework also facilitates individualized dose standardization—different individuals can be assigned configurations achieving equivalent target intensity (e.g., 0.3 V/m) while maximizing focality, ensuring consistent dosing across participants with varying neuroanatomy [35].

For implementation, several strategies are available depending on equipment accessibility. With neuronavigation systems, the coordinates of all five electrodes can be directly localized. Without neuronavigation, electrode holders with the optimized radius can be prefabricated, and the central electrode position and orientation can be manually measured using cranial landmarks. Alternatively, all electrode positions can be converted to CPC coordinates and marked on a customized cap with pre-positioned holes—a convenient solution for repeated sessions and home-based applications. The discretization of radius and orientation parameters represents a practical balance between solution granularity and computational efficiency, though these can be adjusted depending on application needs.

Several limitations warrant consideration. First, our validation was conducted exclusively in healthy young adults. Pathological conditions may alter tissue conductivity profiles, and aging is associated with increased cerebrospinal fluid volume and cortical atrophy [36]. Whether the SGP_MSS_ constraints generalize to patient populations and older adults requires further investigation. Second, 4×1 HD-tES inherently produces electric fields concentrated in superficial cortical regions. For deep brain targets, alternative approaches such as temporal interference stimulation may be more appropriate. Third, our analysis used electrodes with a 5 mm radius. Larger electrode sizes may alter electric field characteristics; future work should evaluate how electrode size interacts with SGP parameters [37]. Fourth, the CPC-based coordinate system does not fully cover certain facial regions, potentially excluding some electrode placements. While this is unlikely to impact most cortical targets, it should be considered for specific applications. Finally, manual electrode placement based on SGP parameters inevitably introduces spatial error. Previous studies have characterized such errors in landmark-based positioning [24], and future investigations should assess how measurement variability affects targeting performance to establish quality control procedures for clinical implementation.

Future research should address several directions to enhance clinical utility. First, validation of whether enhanced targeting precision translates to improved therapeutic outcomes is essential. Controlled studies comparing optimized montages to standard protocols across applications—including depression treatment, motor rehabilitation, and cognitive enhancement—would link computational optimization to functional efficacy. Second, developing population-based targeting strategies is an important priority. While individualized optimization requires MRI scans, probabilistic targeting approaches derived from large-scale optimization datasets could provide group-level optimal montages that accommodate neuroanatomical variation without individual imaging [27]. Third, implementation in home-based therapy settings warrants systematic evaluation of placement accuracy, user adherence, and therapeutic consistency in unsupervised environments to establish guidelines for remote tES applications [38]. Together, these developments would facilitate translating individualized 4×1 HD-tES optimization from research settings to routine clinical practice, extending precision brain stimulation to broader patient populations.

## Conclusions

The SGP-4×1 HD-tES framework reduces computational complexity by over 90% while guaranteeing global optima identification, substantially outperforming conventional 10-10 placement and matching lead-field-free optimization with greater cross-subject stability. Its scalp-based parameterization enables neuronavigation-free electrode positioning, supporting translation to routine clinical and home-based settings.

## Declarations

### Ethics approval and consent to participate

Data collection was approved by the Research Ethics Committee of the Brain Imaging Center at Southwest University (approval number: spy-2012–012; June 3, 2012). All participants provided informed consent.

### Consent for publication

Not applicable.

### Availability of data and materials

The structural MRI data used in this study are from the Southwest University Longitudinal Imaging Multimodal (SLIM) dataset, which is publicly available at https://doi.org/10.1038/sdata.2017.17. High-resolution figures and extended supplementary materials (Figure S2.zip and Figure S3.zip containing individual-level results for all 30 subjects) are available at [https://figshare.com/s/c9c1f53d95c7eb22087d] (will be made public upon acceptance). Source data for all main figures are provided with this paper. Additional simulation data are available from the corresponding author upon reasonable request.

### Competing interests

The authors declare that they have no competing interests.

### Funding

This work was supported by the National Natural Science Foundation of China (Grant Nos. 82071999 and 61431002) and the Lingang Laboratory & National Key Laboratory of Human Factors Engineering Joint Grant (Grant No. LG-TKN-202205-01) awarded to Chaozhe Zhu.

### Authors’ contributions

FL conceived the study, designed the methodology, performed simulations and data analyses, and wrote the initial draft of the manuscript. SL assisted with data processing and experimental validation. KW and YC contributed to electric field simulations and visualization. ZZ and HC contributed to data preprocessing and auxiliary analyses. TC provided technical support and manuscript editing. CZ conceived and supervised the entire study, secured funding, and critically reviewed and revised the manuscript. All authors read and approved the final manuscript.

## Acknowledgements

The authors thank all colleagues in the lab for their helpful discussions and support throughout this study.

## Notes

https://figshare.com/s/c9c1f53d95c7eb22087d

### Competing Interest Statement

The authors have declared no competing interest.

