## Supplementary for "An Efficiency Targeting Parameter Space for Personalized 4×1 HD-tES: Montage Description, Optimization and Application"

1. **MRI Acquisition Parameters**

High-resolution T1-weighted structural MRI scans were acquired using a magnetization-prepared rapid gradient-echo (MPRAGE) sequence with the following parameters: repetition time (TR) = 1,900 ms, echo time (TE) = 2.52 ms, inversion time (TI) = 900 ms, flip angle = 9°, matrix size = 256 × 256, number of slices = 176, slice thickness = 1.0 mm, and voxel size = 1 × 1 × 1 mm³.

2. **Tissue Conductivity Values**

Tissue conductivity values were assigned as follows: white matter (0.126 S/m), gray matter (0.275 S/m), cerebrospinal fluid (1.654 S/m), skull (0.010 S/m), and scalp (0.465 S/m) [1].

3. **Scalp Position Sampling Procedure**

Candidate central electrode positions were first defined on the high-density CPC700 grid, in which both proportional coordinates are uniformly subdivided into 700 steps, generating approximately 490,000 continuous scalp locations. From this dense parameter space, a set of uniformly distributed scalp points was obtained using Fibonacci-sphere resampling, resulting in a practical sampling grid with an effective spacing of approximately 4 mm. This procedure yields approximately 4,500 initial candidate positions with homogeneous coverage across the scalp surface.

4. **Feasible Parameter Space Coverage Analysis**

The number of feasible central electrode (CE) positions varies systematically with both radius (r) and orientation (φ), reflecting geometric constraints imposed by anatomical exclusion zones (Figure S1a, b).

Effect of Radius. As radius increases from 25 mm to 70 mm, the mean number of valid positions decreases from approximately 2,850 to 1,480 across subjects. This reduction occurs because larger radii increase the likelihood of return electrodes falling into periorbital or periauricular exclusion zones.

Effect of Orientation. Within each radius level, φ = 45° consistently yields the highest number of valid positions, while φ = 0° and φ = 75° yield fewer. This pattern likely reflects that diagonally positioned return electrodes more effectively avoid anterior and lateral exclusion zones.

Inter-Subject Variability. At smaller radii (25–35 mm), the standard deviation across subjects is relatively low (60–85), indicating limited impact of individual head geometry. At larger radii (>50 mm), variability increases substantially (85–115), with φ = 0° showing the highest variability at r = 70 mm (STD = 113.5).

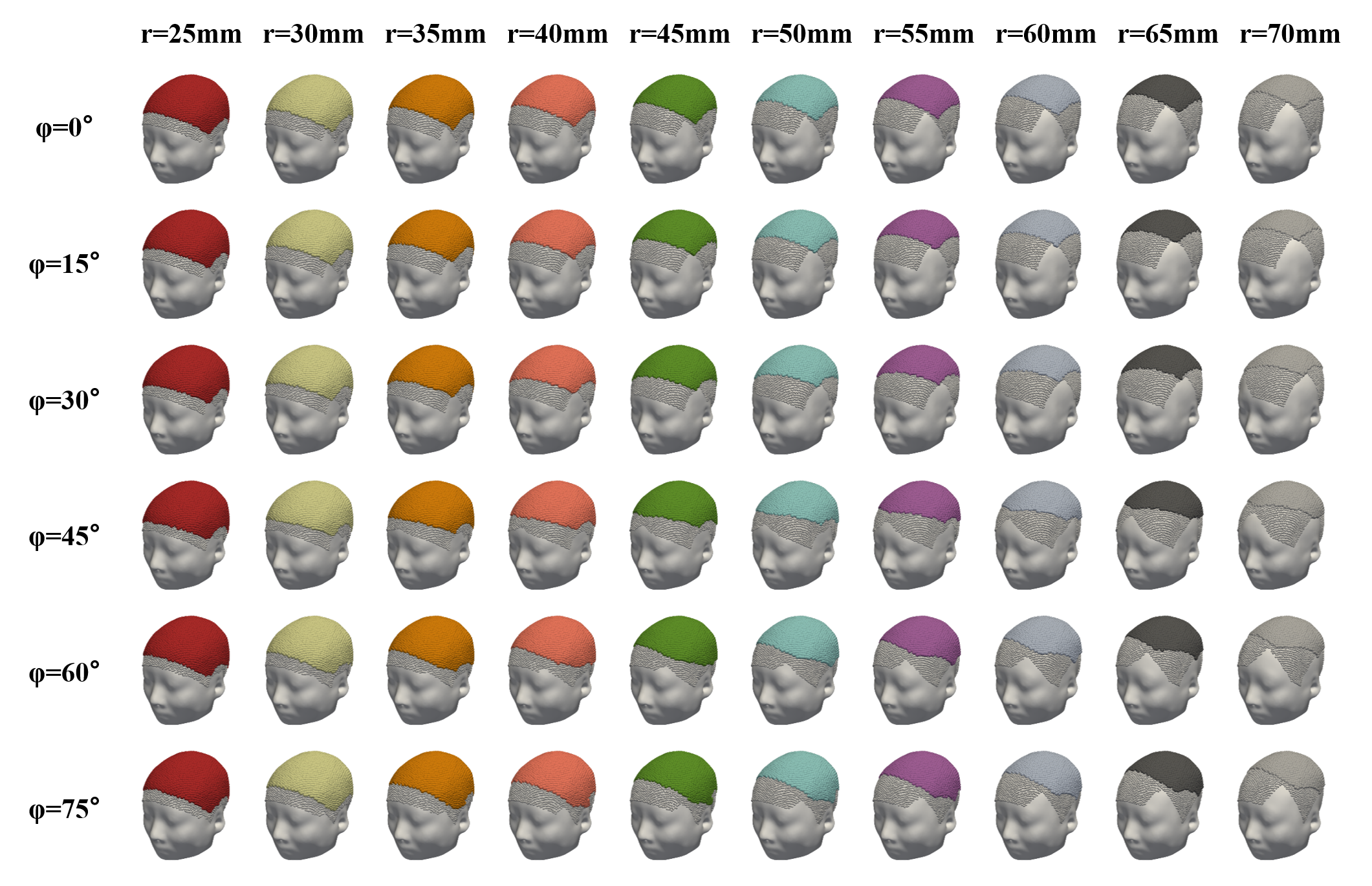

**Figure S1a.** Feasible CE positions for a representative subject across all radius (columns) and orientation (rows) combinations. Colored regions indicate valid placement areas; coverage decreases with increasing radius.

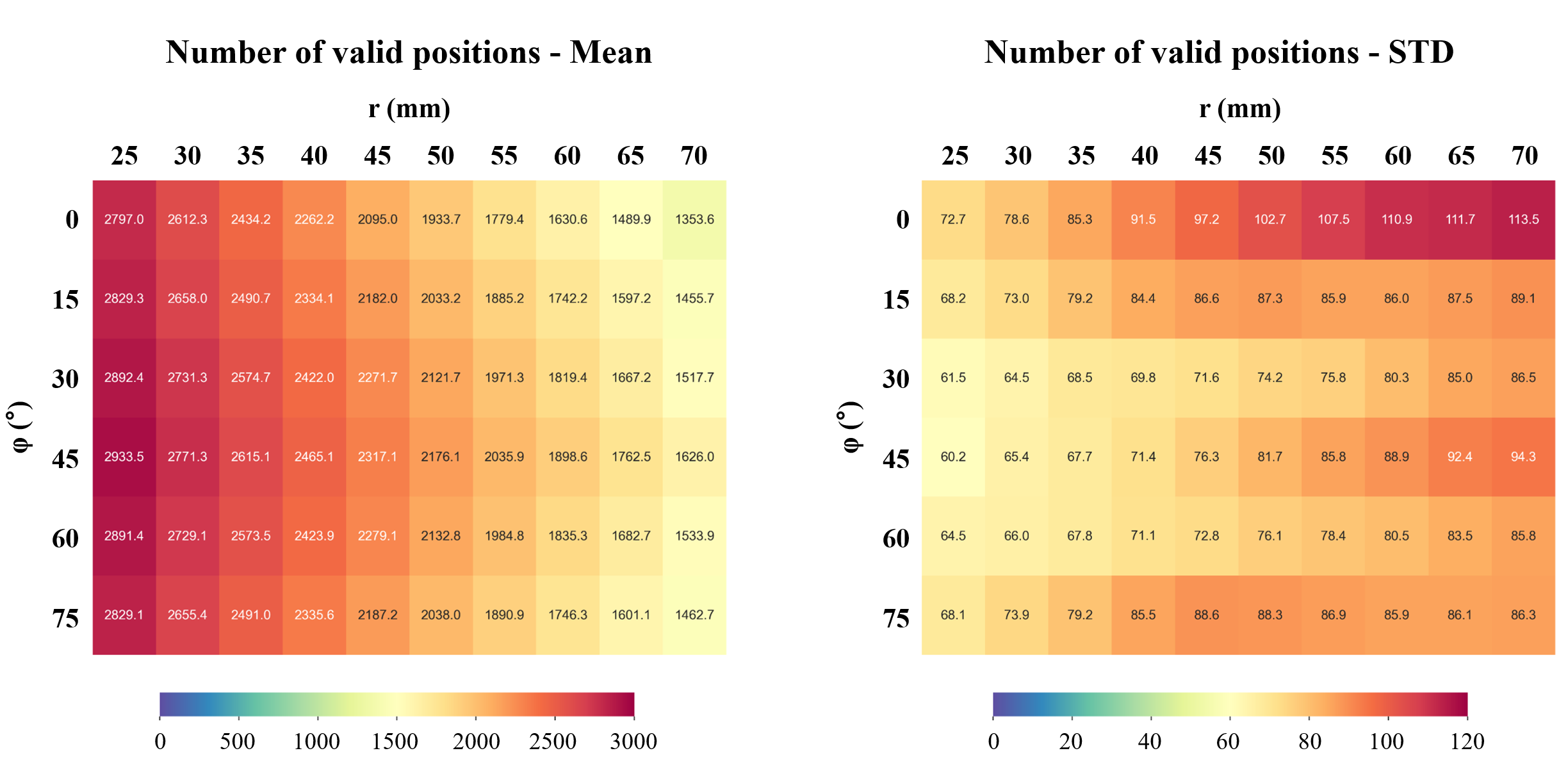

**Figure S1b.** Quantitative summary across 30 subjects. Left: mean number of valid positions. Right: standard deviation reflecting inter-individual variability.

5. **Conventional 10-10 Electrode Configurations**

For the Pareto-based validation, standard 4×1 configurations commonly used in clinical and research settings were applied for each target ROI: C3 served as the central electrode for L_M1 with CP3, FC3, C5, and C1 as return electrodes; F3 served as the central site for L_DLPFC with F5, F1, AF3, and FC3 as returns; P3 was used for L_IPL with P1, P5, CP3, and PO3 as returns; and CP6 was used for R_TPJ with P6, C6, TP8, and CP4 as returns [2,3].

6. **ROC-Based Evaluation Metrics**

The ROC-based evaluation framework implemented in the Lead-Field Free Optimization Framework (LFOF) [4] defines targeting intensity as the proportion of ROI volume where the electric field exceeds a specified threshold (default: 0.3 V/m), and targeting focality as the proportion of gray-matter volume outside the ROI where the field exceeds another threshold (default: 0.1 V/m). These metrics are mapped to ROC space, and the optimal configuration is defined as the point minimizing the Euclidean distance to the ideal point (0, 1).

7. **Lead-Field Simulation for Method Comparison**

Since LFOF employs rapid electric field approximation during optimization and requires a conventional lead-field simulation for the final optimal configuration, the same lead-field simulation procedure was applied to the 10-10 and SGPMSS optimal configurations to ensure comparability across all three methods.

8. **Target-Specific SGPMSS Configuration Details**

The number of configurations within the SGPMSS varied by target location: L_M1 (17,005 ± 1,134 configurations, 13.2 ± 1.3% of full space), L_DLPFC (11,355 ± 809 configurations, 8.8 ± 0.8%), L_IPL (12,812 ± 922 configurations, 9.9 ± 0.7%), and R_TPJ (7,662 ± 1,123 configurations, 5.9 ± 0.8%).

9. **Sensitivity Analysis Across Subjects and ROI Sizes**

To verify the sensitivity of findings presented in Figures 2 and 3 of the main text, exhaustive simulations were performed across all 30 subjects, four representative ROIs (L_M1, L_DLPFC, L_IPL, R_TPJ), and three ROI sizes (5 mm, 10 mm, 15 mm radius spheres). Complete results are provided in two compressed archives: Figure S2.zip (targeting performance distributions) and Figure S3.zip (parameter-performance relationships).

Both archives are organized hierarchically: ROI size → target region → individual subject. Across all conditions, the patterns were consistent with the main text: (1) the characteristic performance envelope and Pareto front structure remained stable; (2) targeting performance decreased systematically with increasing geodesic distance from target; (3) larger radii yielded higher intensity but lower focality; and (4) orientation exerted relatively modest effects. These findings confirm that the results generalize robustly across the full cohort and ROI definitions.

10. **Detailed Statistics for Pareto-Based Validation**

Targeting intensity comparison (matched for focality): L_M1 (0.36 ± 0.08 vs 0.18 ± 0.04 V/m, t = 16.75, p < 0.0001), L_DLPFC (0.41 ± 0.10 vs 0.35 ± 0.07 V/m, t = 8.28, p < 0.0001), L_IPL (0.33 ± 0.08 vs 0.22 ± 0.05 V/m, t = 9.59, p < 0.0001), R_TPJ (0.26 ± 0.05 vs 0.21 ± 0.03 V/m, t = 9.05, p < 0.0001).

Targeting focality comparison (matched for intensity): L_M1 (19.15 ± 2.67 vs 8.48 ± 0.92, t = 23.67, p < 0.0001), L_DLPFC (23.35 ± 4.59 vs 18.91 ± 2.11, t = 7.42, p < 0.0001), L_IPL (20.14 ± 3.08 vs 13.50 ± 1.35, t = 12.24, p < 0.0001), R_TPJ (10.55 ± 0.96 vs 9.05 ± 0.63, t = 8.55, p < 0.0001).

11. **Detailed Statistics for ROC-Based Validation**

ROC distance (mean ± SD) for each method:

L_M1: 10-10 (0.94 ± 0.12), LFOF (0.24 ± 0.28), SGPMSS (0.21 ± 0.24). L_DLPFC: 10-10 (0.20 ± 0.24), LFOF (0.16 ± 0.28), SGPMSS (0.07 ± 0.02). L_IPL: 10-10 (0.83 ± 0.24), LFOF (0.19 ± 0.22), SGPMSS (0.17 ± 0.14). R_TPJ: 10-10 (0.96 ± 0.07), LFOF (0.33 ± 0.24), SGPMSS (0.26 ± 0.11).

Pairwise comparisons: SGPMSS vs 10-10: L_M1 (t = 16.67, p < 0.0001), L_DLPFC (t = 3.02, p = 0.005), L_IPL (t = 14.36, p < 0.0001), R_TPJ (t = 33.21, p < 0.0001). LFOF vs 10-10: L_M1 (t = 13.94, p < 0.0001), L_DLPFC (p = 0.55), L_IPL (t = 12.12, p < 0.0001), R_TPJ (t = 13.77, p < 0.0001). SGPMSS vs LFOF: all p > 0.05.

12. **Validation of SGP_MSS_ Optimization Across Different ROI Sizes**

To assess the robustness of the SGP_MSS_ optimization results presented in Validation 1 (Figure 5 of the main text), we repeated the Pareto-based comparison between SGP_MSS_ optimization and conventional 10–10 placement using two additional ROI sizes: 5 mm and 15 mm radius spheres. The main text reported results for 10 mm ROI size; here we present the complete results across all three ROI definitions.

Results for ROI Size = 5 mm. SGP_MSS_ optimization significantly outperformed the conventional 10–10 placement for all four targets in both targeting intensity and targeting focality (all p < 0.0001; Figure S4). For L_M1 (C3), SGP_MSS_ achieved higher intensity (0.40 ± 0.09 vs. 0.20 ± 0.04 V/m, t = 15.67) and higher focality (22.45 ± 3.91 vs. 9.53 ± 1.24, t = 19.13). For L_DLPFC (F3), intensity improved from 0.40 ± 0.09 to 0.49 ± 0.13 V/m (t = 8.01) and focality from 21.43 ± 2.97 to 28.72 ± 6.91 (t = 7.90). For L_IPL (P3), intensity improved from 0.24 ± 0.06 to 0.37 ± 0.09 V/m (t = 9.69) and focality from 14.82 ± 1.56 to 23.74 ± 3.51 (t = 13.72). For R_TPJ (CP6), intensity improved from 0.19 ± 0.03 to 0.25 ± 0.05 V/m (t = 6.30) and focality from 8.48 ± 0.73 to 10.00 ± 1.29 (t = 5.86).

Results for ROI Size = 15 mm. Similar improvements were observed with the larger ROI size (all p < 0.001; Figure S4). For L_M1 (C3), SGP_MSS_ achieved higher intensity (0.32 ± 0.07 vs. 0.16 ± 0.03 V/m, t = 16.92) and higher focality (15.14 ± 2.07 vs. 7.64 ± 0.80, t = 21.36). For L_DLPFC (F3), intensity improved from 0.31 ± 0.06 to 0.32 ± 0.06 V/m (t = 4.50, p < 0.001) and focality from 16.55 ± 1.62 to 19.10 ± 2.77 (t = 7.62). For L_IPL (P3), intensity improved from 0.19 ± 0.04 to 0.41 ± 0.07 V/m (t = 24.61) and focality from 11.74 ± 1.17 to 17.85 ± 1.64 (t = 22.86). For R_TPJ (CP6), intensity improved from 0.23 ± 0.04 to 0.41 ± 0.06 V/m (t = 39.54) and focality from 9.85 ± 0.80 to 13.30 ± 1.02 (t = 30.44).

These results demonstrate that the performance advantage of SGP_MSS_ optimization over conventional 10–10 placement is robust across different ROI size definitions, confirming the generalizability of the findings reported in the main text.

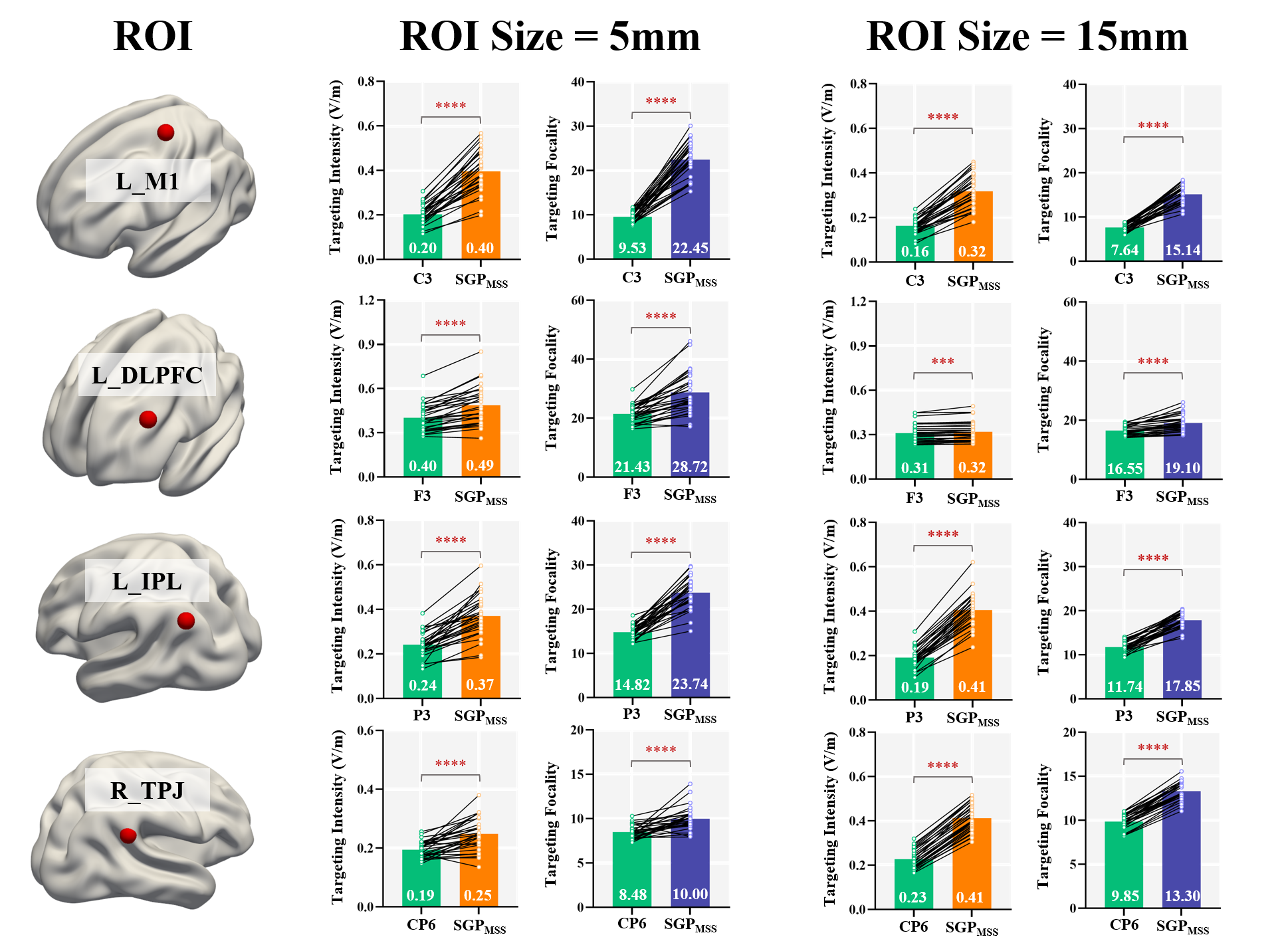

**Figure S4.** Comparison of targeting intensity and targeting focality between 10–10 placement and SGP_MSS_ optimization for ROI sizes of 5 mm (left) and 15 mm (right). Layout follows Figure 5 of the main text. Bar heights represent group means; error bars indicate standard deviation. Black lines connect individual subjects (N = 30). **** p < 0.0001, *** p < 0.001.

13. **Visualization of LFOF Convergence to Suboptimal Plateaus**

As noted in the main text, LFOF occasionally converged to suboptimal solutions for certain subject–ROI combinations. Figure S5 illustrates a representative case for one subject targeting L_IPL.

The left panel displays the optimization trajectory overlaid on the head model. Orange points indicate central electrode positions evaluated during differential evolution, and blue lines trace the search path. The search remained confined to a region far from the target—predominantly around Pz near the vertex—rather than converging toward the scalp above L_IPL. This reflects convergence to a performance plateau where local position variations produce minimal changes in the objective function.

The right panel shows the resulting electric field distribution. Stimulation is concentrated in the superior parietal region, substantially displaced from the intended L_IPL target, directly accounting for the poor ROC-based performance.

This example illustrates a key advantage of SGPMSS: exhaustive enumeration within a physically constrained search space guarantees global optimum identification, avoiding convergence failures inherent to stochastic optimization.

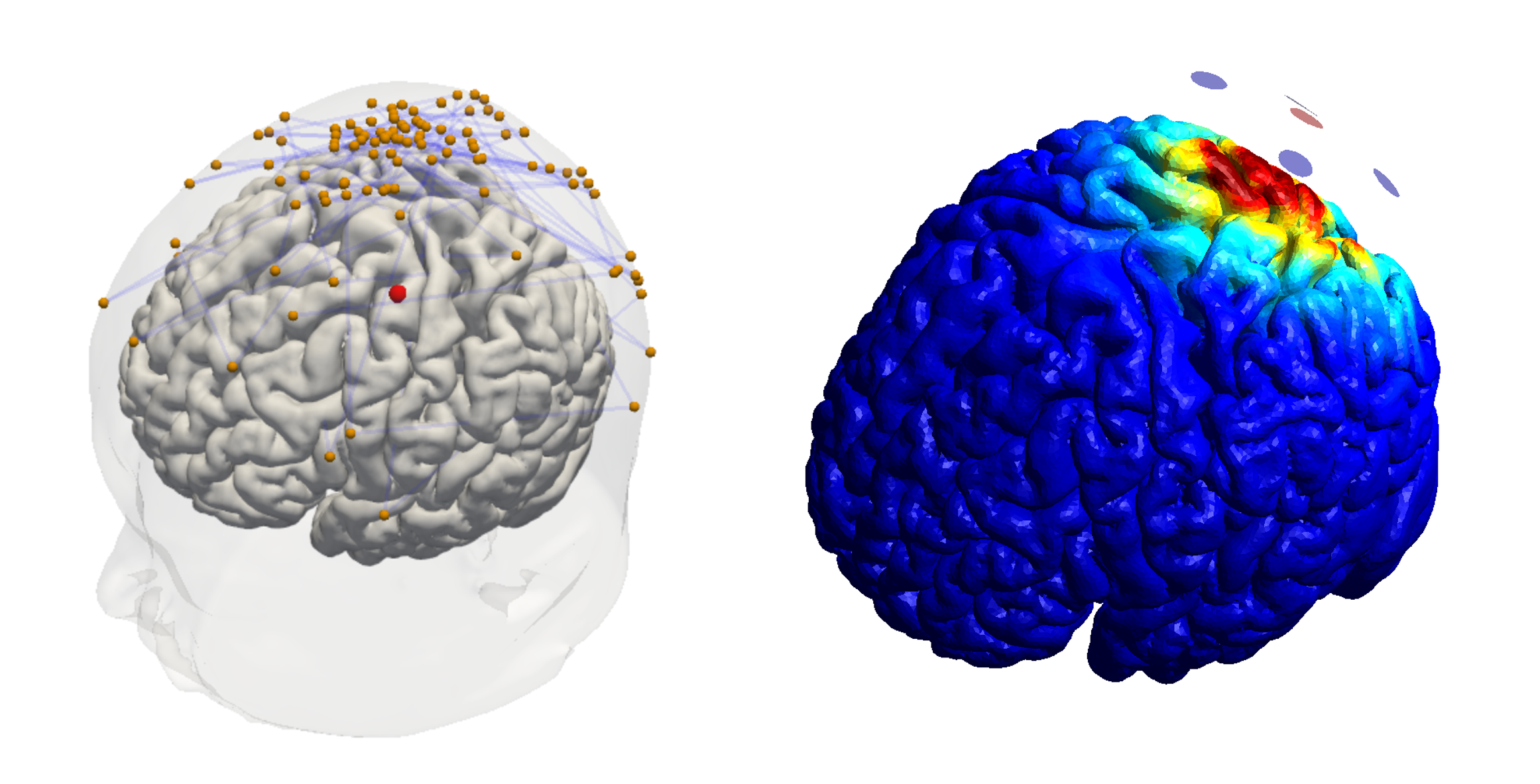

**Figure S5.** Visualization of LFOF convergence to a suboptimal plateau for a representative subject targeting L_IPL. Left: Head model showing the LFOF optimization trajectory (orange points: evaluated central electrode positions; blue lines: search path) and target ROI location (red sphere on cortex). The search remained confined to a region around Pz, far from the optimal placement area above the L_IPL target. Right: Electric field distribution on the cortical surface resulting from the final LFOF-optimized configuration, showing stimulation concentrated near the vertex rather than at the intended target location.
